# Infantile engram modulates memory formation during adulthood

**DOI:** 10.64898/2026.06.19.733374

**Authors:** Joo Hee Yang, Sunhoi So, Jin-Hee Han

## Abstract

Infantile amnesia refers to the inability to recall early-life experiences, despite their lasting influence on adult behavior. Recent evidence suggests that memory traces formed during infancy may persist in a silent engram state. However, whether and how these silent engram cells contribute to adult memory formation remains unclear. To address this, we examined whether experiences during infancy affect adult memory formation using contextual fear conditioning in mice. Consistent with previous studies, we observed infantile amnesia 30 days later. Despite amnesia, these mice exhibited memory enhancement upon retraining as adults, suggesting that silent infantile engrams influence adult memory. Notably, this effect was context-specific. Using a cellular tagging system and ablation approaches, we found that infantile engram cells in the infralimbic cortex (IL) were crucial for this memory enhancement. Overall, these findings demonstrate that infant engram cells in the IL are re-recruited into adult memory traces, providing a neural mechanism by which early-life experiences shape adult memory formation through relearning.

## Main

Early life experiences are known to shape long-term emotional and cognitive trajectories. Infantile amnesia, the inability to recall early childhood memories, may not reflect permanent memory loss, but rather the persistence of partial infant memory traces (Freud, 1905; Josselyn & Frankland, 2012; Rubin, 2000). Instead, accumulating evidence suggests they may persist in a latent, inaccessible state and influence adult behavior unconsciously (Callaghan & Richardson, 2012; Contreras *et al*, 2024; Guskjolen *et al*, 2018; Jacobs & Nadel, 1999; Mineka & Oehlberg, 2008).

Human studies have shown that early childhood memories are not uniformly inaccessible (Akers *et al*, 2012; Peterson, 2002; Rubin, 1982, 2000; Wetzler & Sweeney, 1986). Infantile memory is found to be encoded at the time the event occurred despite their limited accessibility later in life. Supporting this infantile memory formation, young children are capable of recalling early-life experiences over certain periods, indicating that at least some infant memories persist over time (Bauer *et al*, 1994; Bauer *et al*, 1998; Bauer & Wewerka, 1995; Myers *et al*, 1987; Peterson, 2002; Peterson & Whalen, 2001).

Consistent with findings from human studies, animal models have provided critical insights into the neural underpinnings of infant memory trace. In particular, infant memory can be reinstated through behavioral reactivation protocols (Travaglia *et al*, 2016). Infant memories are stored as latent memory traces that persist despite being functionally inaccessible under normal conditions. Supporting this idea, latent infant memory can be reinstated or artificially reactivated through direct stimulation of engram cells. This leads to the concept of “silent engram”, functionally inaccessible yet enduring memory traces (Guskjolen *et al*., 2018; Josselyn & Tonegawa, 2020). Similar to this concept, recent studies have shown that learning experiences during infancy can establish long-lasting memory schemas that subsequently influence behavior in adulthood (Bessières *et al*, 2026; Contreras *et al*., 2024).

However, it remains unclear whether these dormant memory traces play an active role in shaping behavior in adulthood. Using the contextual fear conditioning paradigm in mice, we first found that relearning in adulthood enhanced fear memory. To directly examine whether memory engram cells formed during infancy influenced adult memory, we used an activity-dependent engram-tagging approach. By employing the TRAP2 x Ai14 system to label neurons activated during contextual fear learning in infancy, we found that infantile engram cells, specifically those in the infralimbic cortex (IL), were re-recruited to support memory formed by relearning in adulthood. We used diphtheria toxin A subunit (DTA), a widely used and powerful tool for cell ablation, to eliminate infantile engram cells in the IL and found that such cortical infantile engram cells are essential for early-life experiences to modulate memory formation via later relearning in adulthood. This work contributes to a growing understanding of how early-life experiences become embedded in the brain and later shape cognitive outcomes through persistent, yet silent, memory traces (Guskjolen *et al*., 2018; Josselyn & Tonegawa, 2020).

## Results

### Infant learning enhances memory formed by relearning in adulthood

To confirm that infant mice rapidly forget fear memories, we used wild-type (WT) mice and conducted CFC. We chose CFC because previous studies have confirmed that contextual fear memory lasts for ∼24 hours. Then, infant mice gradually forgot the fear memory (Akers *et al*., 2012; Akers *et al*, 2014). Two groups of mice [postnatal 17 (P17) and adult] were trained for CFC. They were then tested 1 and 30 days later, in the same context as the infant CFC session. The contextual fear memory was measured as freezing levels, the index of the contextual fear memory. In P17 mice, freezing level significantly declined 30 days after CFC (Fig. 1a, b). In contrast, adult mice exhibited intact freezing levels in all retention delays, confirming previous reports of infantile amnesia (Fig. 1c, d).

**Figure 1.**
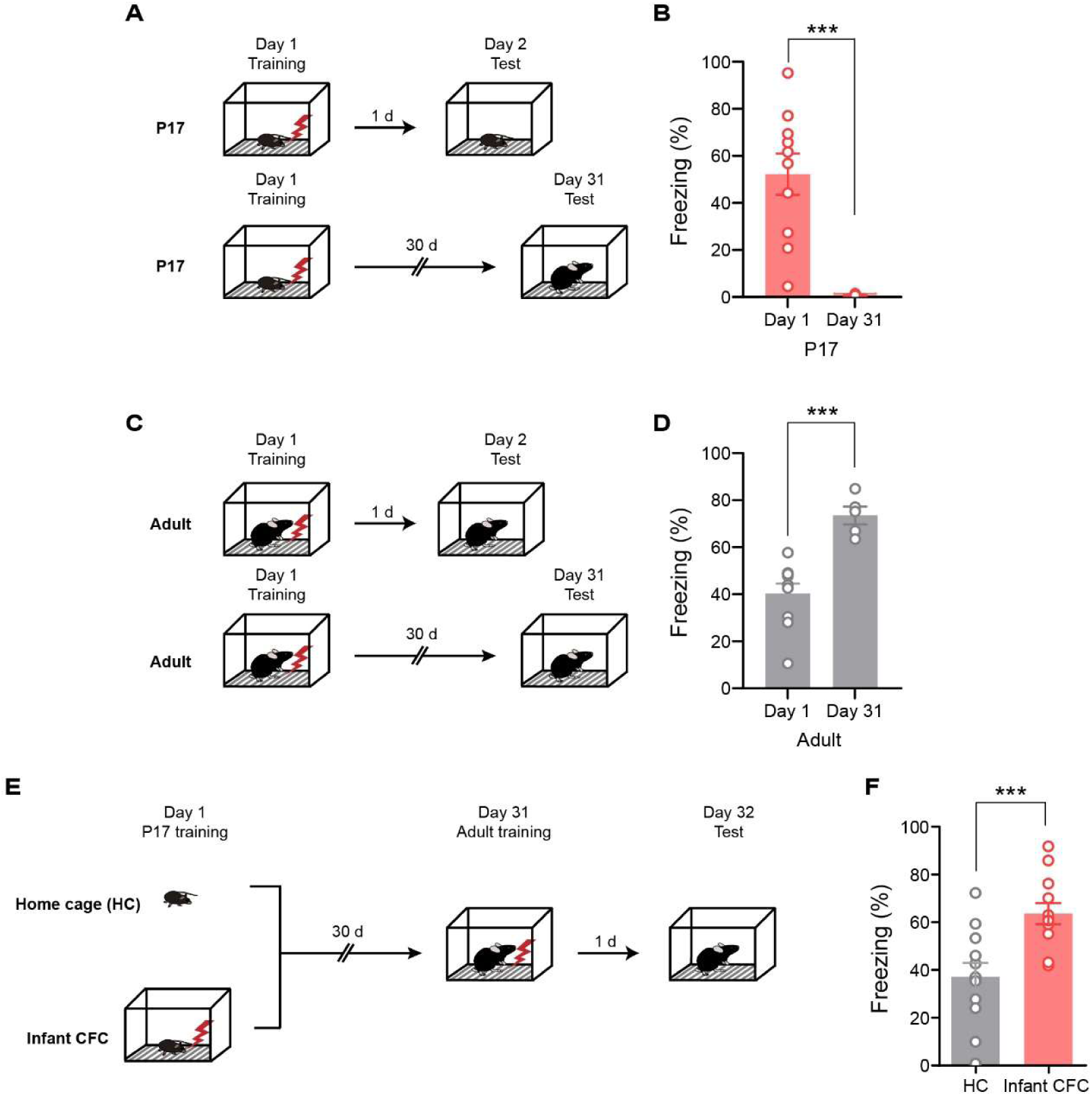
infantile contextual experience enhances adult fear memory expression. **(A, C, and E)** Behavioral paradigm. **(A, C)** Mice were trained for CFC, and contextual test 1 day and 30 days after CFC **(B)** The mean percentage of freezing level during the memory test. P17 mice dramatically decreased freezing levels of Day 1 (n=10 mice) than of Day 31 (n=6 mice, *p=*0.0005, unpaired t-test) after CFC, showed infantile amnesia. (**D)** In contrast, adult mice exhibited intact freezing level after Day 1 (n =10 mice) and Day 31 (n=5 mice, *p=*0.0003, unpaired t-test). (**E)** Behavioral paradigm. In HC group (n=12 mice), P17 mice stayed in HC. 30 days after, they were given CFC, and then 1 day after, they underwent the contextual test. In infant CFC group (n=12 mice), P17 mice were given CFC, and the remaining procedures are same as the HC group. **(F)** The freezing level in infant CFC group was significantly higher than the HC group (*p*=0.0014, unpaired t-test). \*\*\**p* <0.001. Data are mean ± s.e.m

Next, we examined whether the initial contextual fear memory formed in infancy can affect memory formation in adulthood at the behavioral level. There were two groups of P17 mice: the experimental group (Infant CFC) was trained at P17 (Day 1) and retrained at Day 31, whereas the control group (Home cage) was trained only at Day 31 (Fig. 1e). In the contextual memory test, they were returned to the conditioned context and assessed freezing levels. The freezing level in the infant CFC group was significantly higher than that in the control group (Fig. 1f).

To test whether exposure to CFC during infancy affects general locomotor activity or anxiety, P17 mice were trained with CFC and, 30 days later, underwent an open-field test and subsequently an elevated plus-maze test (Fig. EV1a). Mice remained in home cage during infancy were included as controls. We found no significant differences between groups in either test (Fig. EV1b-g). Early childhood fear experiences may alter stress reactivity in adulthood and thereby increase freezing levels. To rule out this possibility, we measured blood corticosterone levels after CFC in adult mice that had undergone one of the following three conditions during infancy: home cage (HC), immediate shock (IS), or infant CFC (Fig. EV2a). We found that IS and infant CFC groups exhibited higher corticosterone levels than those in HC group (Fig. EV2b), suggesting that exposure to stress during infancy increases stress reactivity. Given that only mice that experienced CFC, not IS, during infancy displayed increased freezing during testing following relearning in adulthood, the increase in freezing cannot be due to the increase in stress-hormone response (Fig. 2b). Together, these results demonstrate that infant learning enhances relearning in adulthood. Despite the infantile amnesia, the infant learning experience strengthens behavioral responses by subsequent relearning in adulthood.

**Figure 2.**
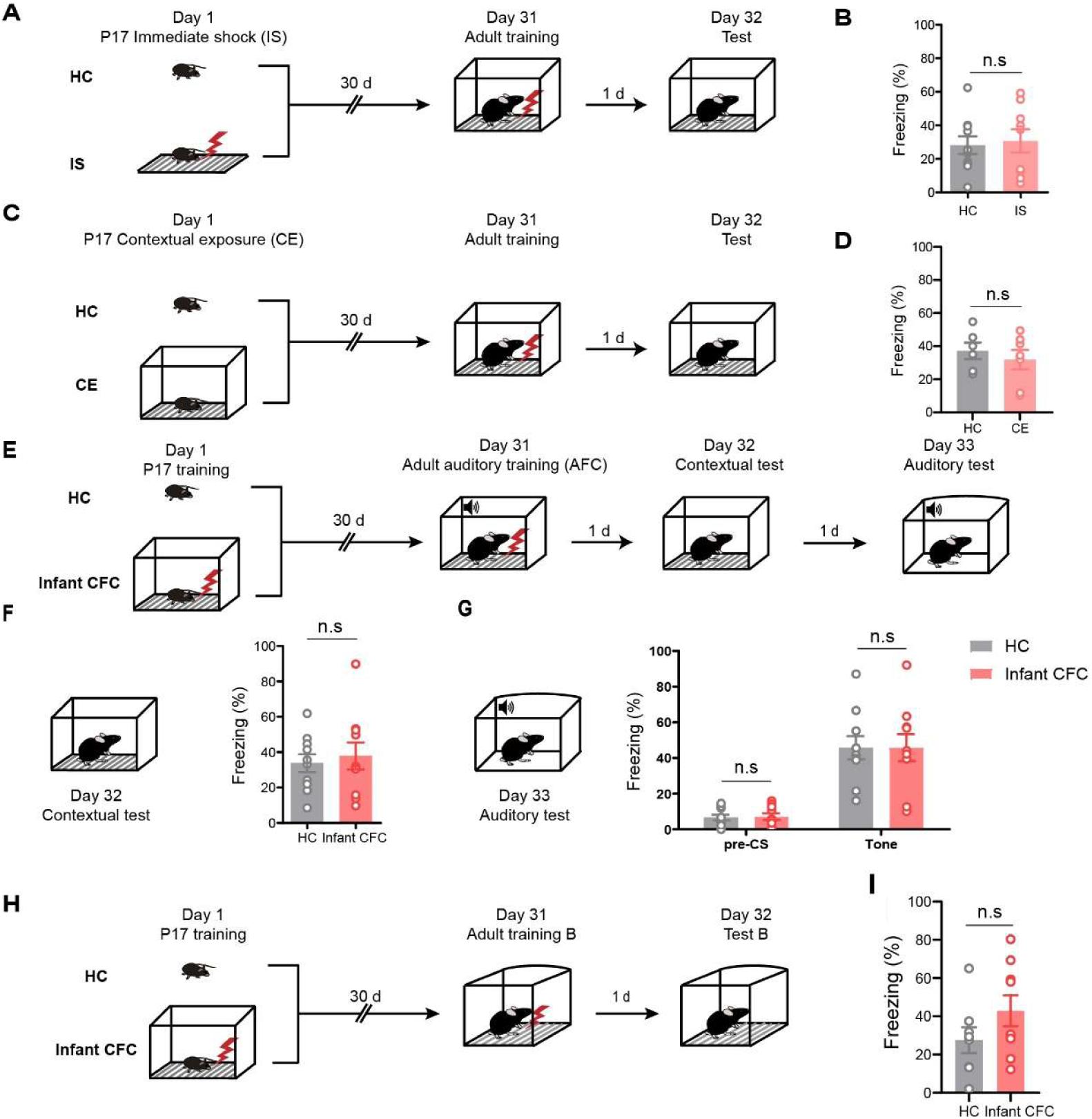
Enhancement of adult fear memory is context-specific. **(A)** Behavioral paradigms. P17 mice stayed HC (n=10 mice) or mice received immediate shock right after the chamber entry (n=9 mice). **(B)** There was no significant difference in freezing level between two groups (*p*=0.7742, unpaired t-test). **(C)** Mice were exposed to the conditioned chamber (n=7 mice) or stayed home cage (n=6 mice). **(D)** There was no significant difference in freezing level between two groups (*p*= 0.5078, unpaired t-test). **(E)** P17 mice underwent contextual fear conditioning (CFC; n = 10 mice) or remained in the home cage (HC; n = 10 mice). 30 days later, all mice were subjected to auditory fear conditioning (AFC). 1 day after AFC, freezing behavior was assessed in the conditioned context, followed by tone-induced freezing. **(F)** In contextual test, there was no significantly difference between them (*p*=0.6603, unpaired t-test). **(G)** In auditory test, there was no significantly difference between them (*p*>0.9999, two-way repeated measures ANOVA with Bonferroni’s multiple comparisons test). **(H)** P17 mice were given CFC (n=8 mice) or stayed HC (n=9 mice). 30 days after, they underwent CFC in Context B, and 1 day after that, they were measured freezing level in Context B. **(I)** There was no significant difference in freezing level between two groups (*p*=0.1735, unpaired t-test). n.s, not significant. Data are mean ± s.e.m

### Adult fear memory enhancement by infant learning is context-specific

We next investigated whether the effect of infant experience on adult memory formation is context-specific. Shock experience or exposure to context during infancy may be sufficient to enhance freezing levels in adulthood. To determine whether the electric shock itself leads to enhanced freezing level in adulthood, we conducted an IS experiment. In the IS group, P17 mice received an electric shock immediately upon entering the context. After 30 days, the mice were trained with CFC. Mice with no IS at P17 were included as a control. 1 day after CFC, all mice from both groups were returned to the conditioned chamber to assess their freezing behavior (Fig. 2a). The freezing behavior observed in the IS was comparable to that of the control group (Fig. 2b). These results indicate that the foot-shock event alone does not enhance memory in adulthood. Moreover, exposure to context did not affect adult memory formation, as we observed no significant difference in freezing levels between groups during the test following CFC in adulthood (Fig. 2c, d).

We next investigated whether infant learning generally affects the formation of fear memories in adulthood. For this purpose, infant mice were trained with CFC and subsequently with auditory fear conditioning (AFC) during adulthood, in which a neutral tone was paired with a shock in the same context as during the infant period. P17 mice underwent CFC and, 30 days later, were trained with AFC in the same context. A group of mice without infant CFC was included as a control. 1 day after the AFC, all groups were tested for contextual fear memory, and 1 day later, for auditory memory (Fig. 2e). The results showed no significant differences between the groups on either the contextual or the auditory memory tests (Fig. 2f, g).

To further demonstrate the context-specificity of infant learning effect on adult memory, we used two distinct contextual conditions. To determine whether memory enhancement observed in adulthood is specific to CFC experienced during infancy, mice were exposed to two distinct fear-conditioning environments (referred to as Context A and Context B). This approach allowed us to assess whether early-life memories exert context-specific effects. P17 mice were trained for CFC in context A and 30 days later, in context B, and tested in context B for contextual fear memory. Mice without infant CFC were included as a control group (Fig. 2h). We observed no significant difference in freezing between groups (Fig. 2i), supporting the context-specific effect of infant learning.

To determine whether the memory enhancement observed in adulthood reflects memory generalization, mice were exposed to two distinct fear-conditioning environments during testing. This approach allows us to assess whether early-life memories generalize across different environments. One group underwent CFC in Context A (infant CFC), while the control group remained in their HC without any fear conditioning. 30 days later, both groups underwent CFC in the context A in adulthood. 1 day after conditioning, all mice were exposed to the conditioned context (context A) to assess freezing behavior as a measure of memory recall. The following day, mice were exposed to a distinct context (context C) to evaluate whether memory enhancement transferred to a novel context (Fig. EV3a). We found a significant difference in freezing between context A and C in both groups (Fig. EV3b). Moreover, all mice showed no significant difference between two groups in discrimination index (DI), indicating successful discrimination between context A and C (Fig. EV3c). The results indicate that the memory enhancement in adulthood by infant learning is not due to a general increase in freezing response and that infant learning does not induce fear memory generalization. Taken together, these results indicate that modulation of adult memory formation by infant learning experiences is context-specific.

### The engram formed in infancy is recruited in adulthood

The observation that infant learning, while forgotten, can influence later memory formation through relearning in adulthood raises the hypothesis that latent, infantile engram cells may be re-recruited to support adult memory. To test this idea, we employed double-transgenic mice, TRAP2 x Ai14, to label cells activated by CFC at P17 and assessed their reactivation during the retrieval of adult memory.

Two groups of TRAP2 x Ai14 mice were trained for CFC at P17. TRAPing occurred immediately after the infant CFC. 30 days after CFC, one group was retrained in the same context as during infancy, while the other group remained in HC. 1 day after, all mice from both groups were returned to the conditioned chamber for memory testing and sacrificed 90 minutes later to determine c-Fos activity (Fig. 3a). As expected, mice in the adult CFC group displayed robust freezing, whereas control mice in the no adult CFC group displayed almost no freezing, indicating infantile amnesia (Fig. 3b). We focused our engram reactivation analysis on the medial prefrontal cortex (mPFC, including the prelimbic, infralimbic, and anterior cingulate cortex) and hippocampus (HPC). Because we found that only very few cells were labeled in the CA1 subfield of the TRAP2 x Ai14 mice, we focused our analysis on CA3 and the dentate gyrus (DG). There was no significant difference between groups in the proportion of c-Fos (+) cells and labeled tdTomato (+) cells in mPFC and HPC (Fig. 3c-e, Fig. EV4). However, the overlap ratio between tdTomato (+) cells (activated during CFC in infancy) and c-Fos (+) cells (activated during memory recall in adulthood) was significantly higher, specifically in the infralimbic cortex (IL), in the adult CFC group compared to the control (Fig. 3f). Comparison with chance level also showed the same result (Fig. 3g).

**Figure 3.**
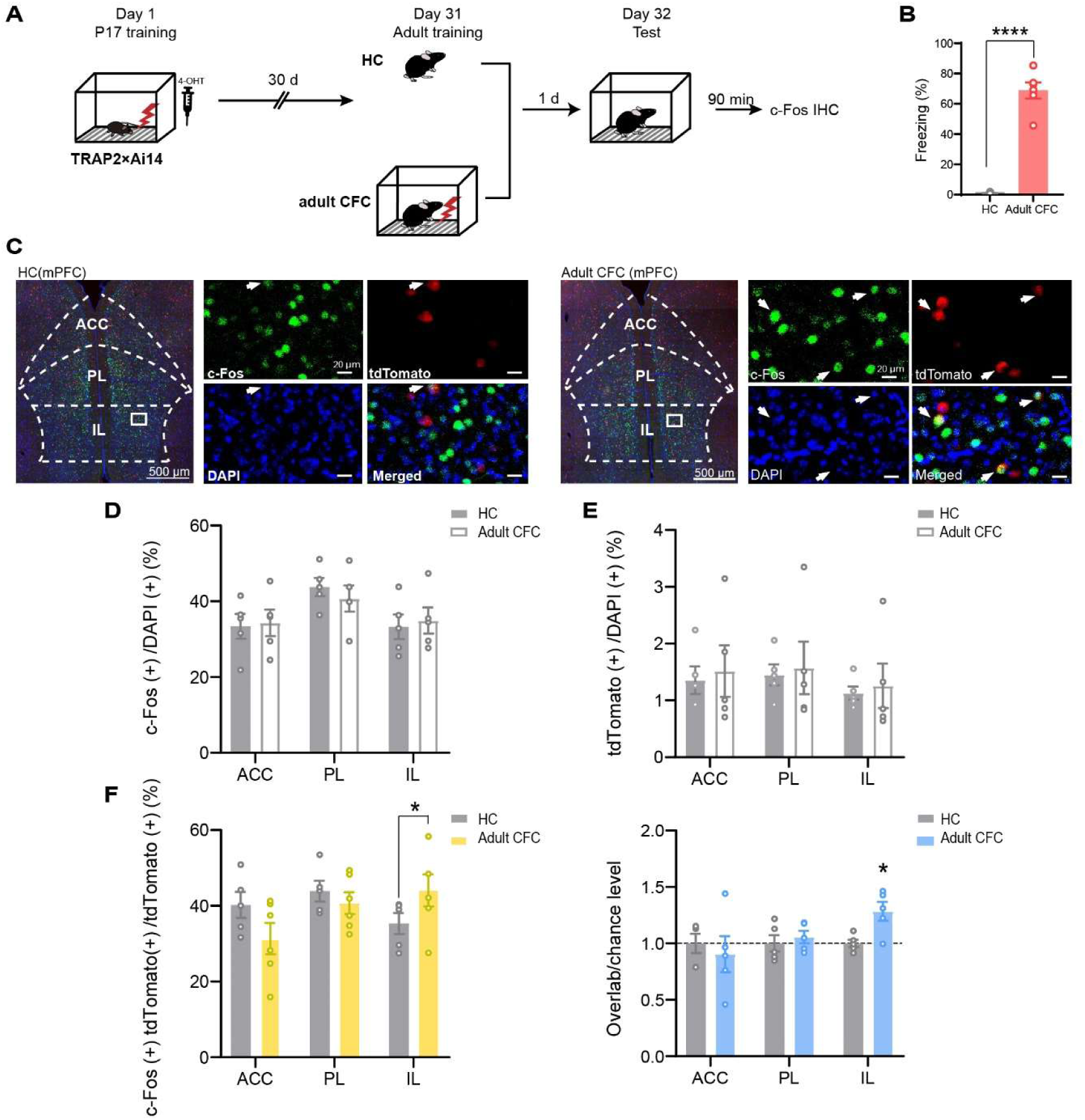
Infant-formed engrams are recruited in the infralimbic cortex (IL) in adulthood. **(A)** Behavioral paradigm. Mice were trained for CFC, followed by 4-OHT injection. 30 days after, they were given CFC (n=4 mice), or remained in HC (n=4 mice) and 90 min after c-Fos immunohistochemistry. **(B)** There was significant difference in freezing level between two groups (n=4 mice, *p* <0.0001, unpaired t-test). **(C)** Representative confocal microscopic images showing cFos+ cells and tdTomato+ cells and their overlap in the IL. Scale bar: 500 μm and 20 μm (inset). (**D,E)** There were no significant differences in cFos (+) cells or tdTomato (+) cells in ACC, PL, IL **(F)** There was significant difference between the overlap in IL (*p*=0.0231, unpaired t-test). **(G)** When the overlap was normalized to chance level, the IL exhibited a significant difference above the hypothetical value of zero (*p*=0.0136, one-sample t-test). \**p*<0.05, \*\*\*\**p*<0.0001. Data are mean ± s.e.m

These results suggest that infantile engram cells in the IL of the mPFC are re-recruited to support adult memory, implying a potential role for these cells in mediating the influence of early-life experiences on adult memory formation.

### Infantile memory engram in IL contributes to adult memory formation

We investigated whether neurons activated during CFC in infancy influence memory formed in adulthood. To test this, we utilized a viral vector encoding diphtheria toxin subunit A (DTA), which induces cell death by disrupting protein synthesis within transduced cells. This enables us to selectively eliminate neurons in the target brain region, particularly in IL, and assess how this affects memory enhancement in adulthood. Mice were injected with AAV-Ef1α-mCherry-DIO-DTA virus (or AAV-Ef1α-mCherry virus as a control) into the bilateral IL of TRAP2 mice on postnatal day 2 (P2) (Fig. 4a). In the presence of 4-OHT, only the neurons that were activated by CFC in infancy express DTA and are thereby selectively ablated. All mice underwent CFC at P17, followed by 4-OHT injection. 30 days later, these mice were retrained and tested the next day in the conditioned context (Context A), then in a novel context (Context C) the following day to assess context specificity (Fig. 4b). Freezing behavior was compared between mice with IL neuron ablation and their respective controls (non-ablated). The ablation group displayed a significantly lower freezing than the control group (Fig. 4c). However, there was no significant difference in discrimination indices between two groups, indicating successful discrimination between context A and C. (Fig. 4d). The number of mCherry (+) cells was significantly reduced in the DTA group compared to the control group, indicating that cells labeled in the IL during infancy were effectively ablated. (Fig. 4e, f). These findings demonstrate that infantile memory traces retained within the IL are re-recruited in the adult engram and contribute to enhance relearning.

**Figure 4.**
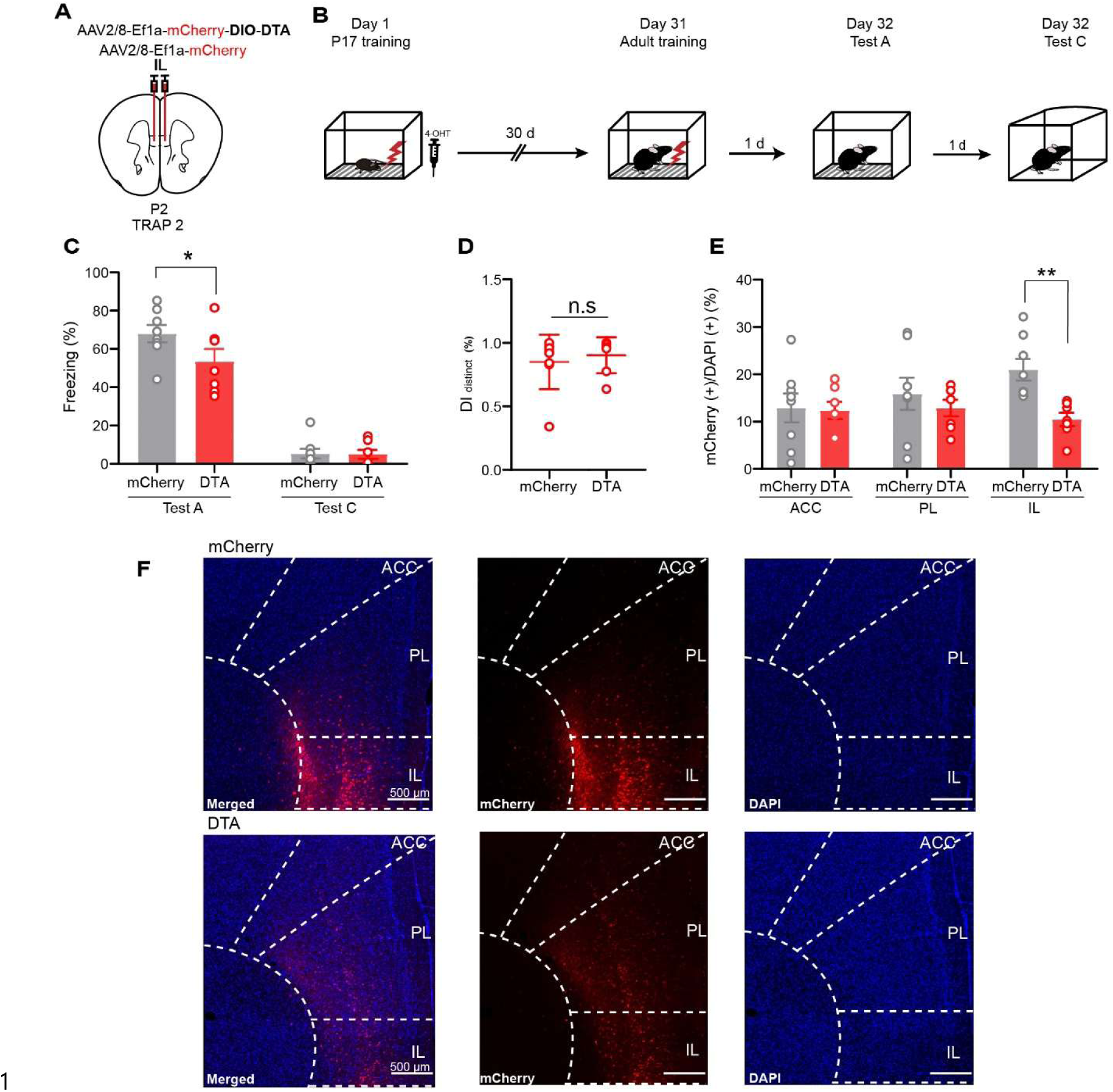
Infantile memory engram in IL contribute to adult memory formation. **(A)** Schematic of viral injection into the IL of TRAP2 mice to label activated neurons in the IL. **(B)** Behavioral paradigm. Mice were trained for CFC, followed by 4-OHT injection. 30 days later, mice underwent contextual fear conditioning (CFC) again (n=7 mice). Freezing levels were assessed 1 day later in the conditioned context (test A), followed by a context-shift test (test C) on the subsequent day. **(C)** There was significant difference between mCherry and DTA groups in freezing level (*p*=0.0230, two-way repeated-measures ANOVA with Fisher’s multiple comparisons test). **(D)** There was no significant different between two groups in discrimination index (DI, *p*=0.5881, unpaired t-test). **(E)** The DTA group showed a significant reduction in mCherry (+) cells compared to the control group in IL (*p*=0.0047, two-way repeated-measures ANOVA with Fisher’s multiple comparisons test). **(F)** Representative images showing mCherry (+) cells in the IL. mCherry expression confirmed on-target virus injection in the IL. Scale bar: 500 μm. n.s, not significant. \**p*< 0.05, \*\**p*< 0.01. Data are mean ± s.e.m

## Discussion

In the current study, we demonstrated that aversive experiences during infancy, while forgotten, enhance re-learning in adulthood, and that the cells activated in the infralimbic region of the mPFC during infancy are crucial for the enhancing effect of early life experiences on memory formation in adulthood. Our findings support the notion that, despite apparent behavioral amnesia, early experiences are not lost but instead leave physical traces in the mPFC in a latent state that contributes to learning in adulthood.

Notably, this enhancing effect was context-specific and not due to a general increase in sensitivity to shock or stress; animals exhibited an enhanced freezing response to the context only when re-experiencing shock in the same context. Given that the freezing response was highly specific to the conditioned context, infantile shock experience did not induce fear memory generalization in adulthood. Moreover, experiencing shock alone during infancy did not affect adult learning, whereas it did increase the stress-hormone response to shock. These findings highlight the highly specific impact of early life experiences on learning in adulthood.

Our findings are consistent with previous reports showing that memories formed in infancy, while rapidly forgotten, can be reinstated later in life and affect learning in adulthood (Bessières *et al*., 2026; Contreras *et al*., 2024; Guskjolen *et al*., 2018; Travaglia *et al*., 2016). As we identified in this study, evidence from recent rodent studies has pointed to the mPFC as a critical brain region that links infantile experiences to adult learning (Bessières *et al*., 2026; Contreras *et al*., 2024). Using the engram tagging technique, a recent work showed that infantile engram cells in the mPFC and their projections to the hippocampus are critical for the facilitating effect of infantile experiences on adult learning (Bessières *et al*., 2026). Together with these findings, our results reinforce the idea that infantile engram cells in the mPFC are re-engaged to enhance learning in adulthood.

We observed no overall difference in c-Fos activity between adult CFC and home cage groups across any of the brain regions we analyzed. Moreover, we found an increase in c-Fos activity in infant-tagged cells, specifically in the IL region of the mPFC, during adult memory retrieval relative to control mice. These observations suggest that infantile engram cells formed in the IL may be re-engaged to participate in the adult engram. This IL-specific reactivation of infant-tagged engram cells is somewhat inconsistent with previous reports emphasizing the importance of the ACC and PL regions of the mPFC (Bessières *et al*., 2026; Contreras *et al*., 2024). This inconsistency may stem from different types of memory used (associative fear memory versus spatial memory) and different memory stages examined for c-Fos activity (encoding versus retrieval).

It is unclear how IL infantile engram cells contribute to memory enhancement in adulthood. The IL is known to be critical for controlling the expression of fear extinction memory in adult mice (Kim *et al*, 2016; Milad & Quirk, 2002). Thus, it is possible that IL infantile engram cells may control the level of fear memory expression in adulthood via an inhibitory role. Alternatively, the IL cells recruited during infant CFC learning may be a critical component of a schema built from early experiences during infancy that promotes congruent learning later in adulthood, as suggested in recent studies (Bessières *et al*., 2026; Contreras *et al*., 2024; Tse *et al*, 2007). Our study is limited by the lack of temporal specificity in manipulating IL engram cells, as we used a DTA-based cell ablation technique. Future studies should use techniques with better temporal resolution, such as optogenetics, to elucidate the precise role of IL infantile engram cells in adult memory formation and retrieval. In addition, because it was technically unfeasible to specifically target viral injection to the IL region at P2 surgery, we cannot rule out that cell ablation in our study was not restricted to the IL but spread to neighboring areas, such as PL.

In conclusion, the findings in this study suggest that early-life memories are stored in a latent state within the mPFC, particularly in the IL-centered circuits, potentially serving as a foundational schema for future learning. While further research using high-temporal-resolution tools such as optogenetics is needed to analyze the precise inhibitory or modulatory dynamics of these cells, our findings reinforce the mPFC’s status as a critical hub linking early-life ontogeny to adult cognitive plasticity. These findings highlight the importance of future research aimed at identifying therapeutic approaches for mental disorders associated with early-life trauma by targeting infantile fear memories that persist in a dormant state into adulthood.

## Methods

### Animals

C57BL/6J mice (P17 days old) were used. The day of birth was designated postnatal day 0 (P0). At 21 days after the birth, infant mice were weaned from their parents, and same sex mice were group-housed (2-5 mice per cage). They were housed in a 12-h light/dark cycle environment at a constant temperature of 22±1°C with 40–60 % humidity. TRAP2 (JAX #030323) mice (+/+) were bred with Ai14 (JAX #007914) mice (+/+) to generate double-transgenic TRAP2 × Ai14 (+/+) mice. For the DTA experiments, TRAP2 (+/+) mice were crossed with C57BL/6J wild-type mice, and offspring heterozygous for the TRAP2 transgene (+/−) were selected for the experiment. Both female and male mice were used, as no behavioral influence of gender was observed. Mice were gently handled for 3 consecutive days for 5 min before behavioral procedures. Food and water were available *ad libitum*. All procedures were approved by the Animal Ethics Committee at the Korea Advanced Institute of Science and Technology (KAIST).

### Virus production

The AAV virus was prepared as previously described (Kwon *et al*, 2014). DNA plasmids (AAV2/8-Ef1a-mCherry-DIO-DTA, AAV2/8-Ef1a-mCherry) were amplified and purified by using a maxi-prep kit (Qiagen #12163). The purified DNA plasmids and viral vector (pAdΔF6 and pAAV2/8) were co-transfected into HEK293T cells by calcium phosphate precipitation. Cells were harvested 72 hours after transfection and the viruses were purified by iodixanol gradient ultracentrifugation. The viral titer was measured by quantitative PCR, and the final titers were: 8.82×10^12^vg/mL for AAV2/8-EF1a-mCherry-DIO-DTA, 5.51×10^12^vg/mL for AAV2/8-EF1a-mCherry.

### Surgery

Postnatal day 2 (P2) TRAP2 (+/−) mice were anesthetized by hypothermia, incubated on ice for 5-10 minutes until the reflex and respiratory rate were lost. After hypothermia, the mice were placed onto a neonatal stereotaxic apparatus equipped with a hand-made head tray. Stereotaxic coordinates for the target structure were determined the junction of the middle of eyes and lateral sinuses over the cerebellum. The injection coordinates were (IL): anterior-posterior (AP): +1.6 mm; medial-lateral (ML): ±0.5mm; dorsal-ventral (DV): −0.8 mm. Instead of the craniotomy, the small burr hole was made using a sterile 27-gauge needle. The viral constructs were delivered via intracranial injection using a pull glass capillary connected to a Nanoject Ⅲ systems (Drummond). The injection volume was 50 nL/s, and the pipette was left in the intended injection place for an additional 1 min to allow diffusion. After 1 min for the diffusion, the pipette was retracted to half of the dorsoventral (DV) depth, maintained in position for 30 s, and then slowly withdrawn. To reduce dam stress, approximately half of pups from each litter were removed for the surgery and the remaining pups were left undisturbed with the dam; after recovery and return of the first half, the second half were processed.

### Drug preparation

4-hydroxytamoxifen (Sigma, Cat# H6278, 50mg) was first dissolved at a concentration of 20 mg/mL in ethanol by shaking at 37 °C for 15 min. The dissolved solution was then aliquoted and stored at −20 °C for up to several weeks. Before injection, 4-OHT was redissolved in ethanol by shaking at 37 °C for 15 min, Chen Oil [a 1:4 mixture of castor oil:sunflower seed oil] was then added to reach a final concentration of 10 mg/mL 4-OHT. The ethanol was evaporated using a vacuum centrifugation and the final 10 mg/mL 4-OHT solutions were stored at 4 °C before use.

### Context fear conditioning

Mice, because they were group-houses, were transferred individually into a fear-conditioning chamber, using a standard mouse cage. The fear-conditioning chamber consists of shock grids available to deliver foot shock and a video camera to monitor the behavior of mice. Video recording and shock manipulation were operated by FreezeFrame5.0 software (Coulbourn Instruments). After entry to the fear-conditioning chamber, mice were able to explore the chamber for 2 min. After 2 min, 5 times foot shock (0.5mA, 2 s) was delivered 120 s, 180 s, 240 s, 300 s and 360 s to the subject for association of context and shock. 1 min after the final foot shock delivery, mice were immediately removed from the fear-conditioning chamber to be returned to the home cage. For repeated contextual fear conditioning, the same procedure was given 30 days after the initial contextual fear conditioning. For context memory test, mice were tested 1 day after the repeated contextual fear conditioning. Mice re-entered the same fear-conditioned chamber. Freezing levels of 5 min after entry were automatically measured as an index of contextual fear memory.

### Immediate shock (IS) experiment

Mice were entered the conditioning context. Immediately after the entry, mice were given a single shock (0.5mA, 2s). Right after shock delivery, mice were returned to the home cage.

### Auditory fear conditioning

Mice were placed in the conditioning chamber (Coulbourn Instruments, PA, USA). After 2 min of exploration, mice received 5 tone–shock pairings (30 s tone, 2.8 kHz, 90 dB), each co-terminating with a 2 s foot shock (0.5 mA). Following an additional 60 s in the chamber, mice were returned to their home cages. 1 day later, contextual memory was assessed for 5 min. 1 day after the contextual test, a tone recall test was conducted in a context-shifted chamber with an acrylic floor and semi-circular walls. After a 2 min pre-CS period, the conditioned tone was presented for a total of 3 min.

### Context exposure experiment

Mice were placed in the conditioning context and allowed to explore freely for 7min without foot shock. After the exploration, they were returned to the home cage.

### Context-shift experiment

#### Context B

Mice were placed into a conditioned chamber, consisted of a semicircular wall and staggered metal grid floor (Med Associates). Mint extract (McCormick & Company) was used as an olfactory cue to provide a distinct environmental context. Mice were allowed to explore freely for 2 minutes and given 5 pairings of a 2-s, 0.5 mA foot shock, with an intershock interval (ISI) of 60 s.

#### Context C

The chamber consisted of a semicircular wall and white acryl floor. Mint extract (McCormick & Company) was used as an olfactory cue to provide a distinct environmental context.

### Open field text

Locomotor activity was assessed in an open field arena (45 × 45 × 25 cm). Mice were placed in the center of the arena and allowed to explore freely for 30 min. The total distance traveled and the number of entries into the center zone (22.5 × 22.5 cm) during the first 20 min were quantified using EthoVision XT (Noldus).

### Elevated plus maze

Anxiety-like behavior was assessed using an elevated plus maze (EPM). The apparatus consisted of two open arms (30 × 5 cm) and two closed arms (30 × 5 × 15 cm) made of white opaque Plexiglas, arranged such that identical arms were positioned opposite each other. The maze was elevated 50 cm above the floor. Mice were placed on the central platform facing a closed arm and allowed to explore for 5 min. The percentage of time spent in the open and closed arms was quantified using EthoVision XT (Noldus).

### Corticosterone assay

Blood samples were collected between 9:00 a.m. and 3:00 p.m., when corticosterone levels are at their nadir in the circadian cycle. Mice were first anesthetized with an intraperitoneal (i.p.) injection of pentobarbital (83 mg/kg of body weight). After the anesthesia, blood was collected from the retro-orbital sinus behind the eye, using heparinized capillary tubes. Samples were immediately transferred to pre-chilled tubes containing lithium (BD Microtainer) and kept on ice. Blood samples were centrifuged about 16000 × g for 10 min at 4 °C. The plasma supernatant was collected and stored at −80 °C for 1 day (Gong *et al*, 2015; Weinert *et al*, 1994).

For analysis, plasma was diluted 1:40 in assay buffer and corticosterone levels were measured using a Corticosterone ELISA kit (Enzo Life Sciences, ADI-901-097) according to the manufacturer’s instructions.

### Immunohistochemistry (IHC)

Brain sections were transferred to 12-well plates and rinsed with PBS. Sections were incubated in a blocking solution (2% goat serum, 0.1% BSA, 0.2% Triton X-100 in PBS) for 1 hour at room temperature and overnight with primary antibodies (rabbit anti-c-Fos antibodies (1:2000; 2250S, Cell Signaling)). After primary antibody incubation, sections were washed in PBS again and incubated with secondary antibodies for 2 hours at room temperature (Alexa Fluor 647 goat anti-rabbit (1:2000; A-21244, Molecular Probes)). Sections were subsequently washed in PBS, mounted on gelatin-coated slides, and coverslipped with DAPI mounting medium (H-1200 or H-2000, Vector Laboratories). Images were obtained using a Zeiss LSM780 confocal microscope.

### Histology

For histological verification of viral expression, brain tissues were collected. Mice were perfused transcardially with PBS followed by 4% paraformaldehyde (PFA). Brains were extracted and post-fixed overnight in 4% PFA at 4 °C. Coronal brain sections (40 μm) were obtained using a vibratome (VT-1200S; Leica). Sections were examined under a fluorescence microscope (Eclipse 80i; Nikon) with reference to the Allen Mouse Brain Atlas. Animals showing insufficient viral expression was excluded from further analysis.

### Cell counting analysis

Cell quantification was performed using ImageJ (FIJI). Brain sections were imaged using a Zeiss LSM780 confocal microscope. Images were acquired using the following objectives: 10× (overview) and 20× objectives for cellular resolution imaging. Fluorophores were excited using standard laser lines. Emission signals were collected using appropriate band-pass filters specific to each fluorophore. Confocal images were acquired identical laser power, gain, and offset settings across all sections to allow quantitative comparison. For each brain region, each z-stack image per brain sections combined using maximum-intensity projection, and converted to 8-bit grayscale. Regions of interest (ROI) were defined according to anatomical boundaries based on the Allen Mouse Brain Atlas. DAPI (+) cells were used to measure as the total number of cells. c-Fos (+) and tdTomato (+) cells were identified, using threshold settings and manually verified. Colocalized cells were manually counted when clear overlap of c-Fos and tdTomato signals was observed within the same DAPI-labeled nucleus. All quantification was performed bilaterally from 3-5 sections per animal.

### Statistical and Behavioral Analysis

all statistical analyses were used GraphPad Prism 10. All tests were two-tailed. For comparisons between two groups, unpaired t-tests were used as appropriate. For comparisons among more than two groups, one-way ANOVA or two-way repeated-measures ANOVA was performed. When ANOVA results were significant, post hoc analyses were conducted using Bonferroni’s or Fisher’s LSD multiple-comparisons tests, as appropriate. Statistical significance was set at \**p* < 0.05, \*\**p*< 0.01, \*\*\**p*< 0.001, and \*\*\*\**p*< 0.0001.

## Acknowledgements

We thank all lab members of Jin-Hee Han’s laboratory for their helpful and constructive discussions and suggestions. We thank the KAIST Bio-Core Center for the use of confocal microscopes and for assistance with imaging experiments.

## Author contributions

Conceptualization: JHY, J-HH

Data acquisition: JHY

Data analyses: JHY, J-HH

Virus production: SS

Visualization: JHY, J-HH

Funding acquisition: J-HH

Supervision: J-HH

Writing – original draft: JHY, J-HH

Writing – review & editing: JHY, SS, J-HH

## Expanded View Figures

**EV 1.**
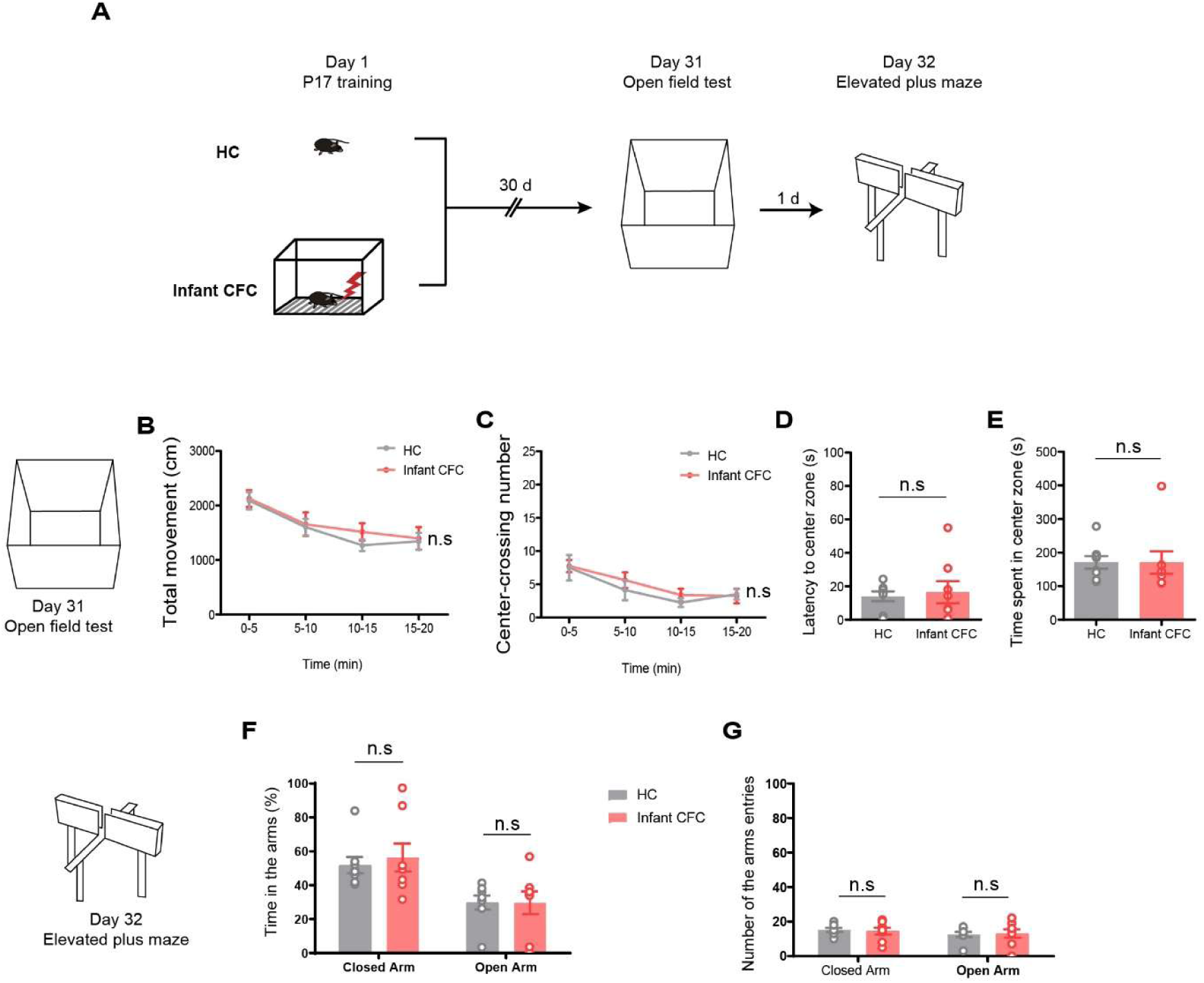
Adult fear memory expression is not enhanced by locomotion or anxiety. **(A)** Behavioral paradigm for open field test (n=8 mice) and elevated plus maze (n=8 mice). Mice underwent CFC or remained in home cage. 30 days after CFC, the open field test was conducted, followed by the elevated plus maze. **(B-E)** Open field test. We found no significant differences in total movement (B), center-crossing numbers (C), latency to center zone (D), and time spent in center zone (E) between two groups. **(F-G)**, Elevated plus maze. There was not significantly difference in time spent (F) and number of the arms entries (G) of open arms and closed arms (*P*>0.05, two-way repeated measures ANOVA with Bonferroni’s multiple comparisons test, unpaired t-test). n.s, not significant. Data are mean ± s.e.m

**EV 2.**
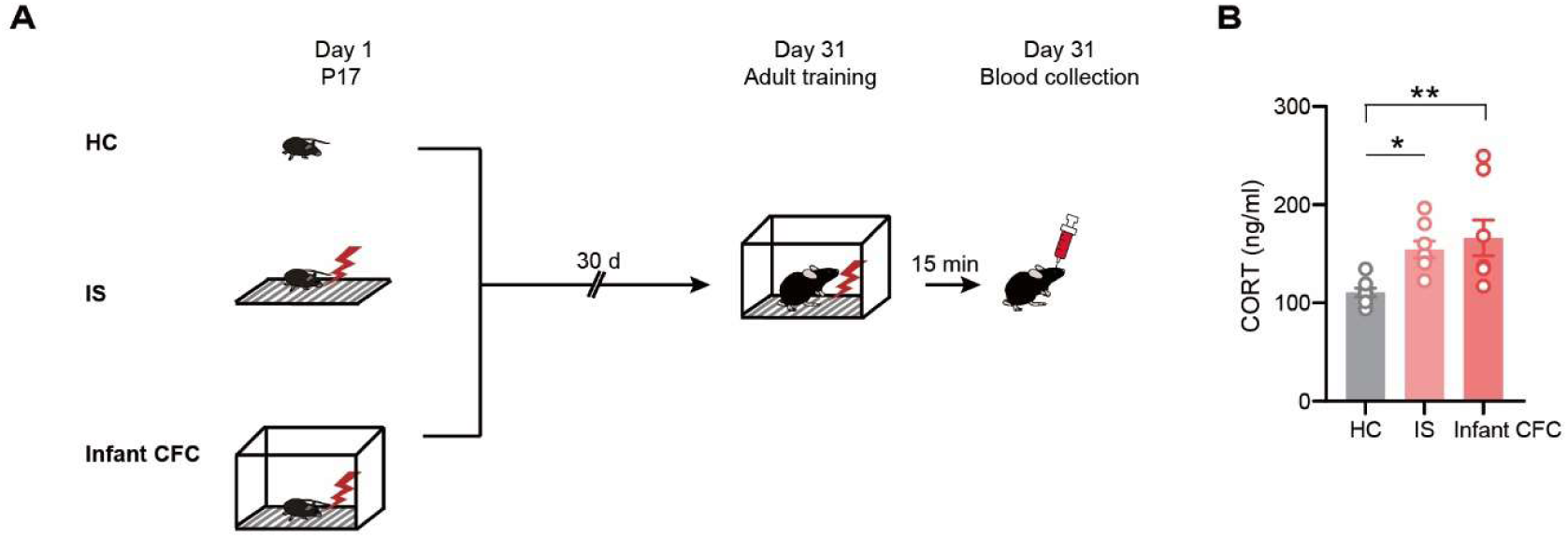
Stress hormone does not lead to the memory enhancement in adulthood. **(A)** Behavioral paradigm for stress hormone experiment (n=8 mice per group). Mice stayed in home cage, were given immediate shock (IS), or underwent CFC. 30 days after, they underwent CFC, and blood was collected 15 min afterward. **(B),** There was significant difference between IS and HC (*p*=0.0423), and infant CFC and HC (*p*=0.0090, ordinary one-way ANOVA with Tukey’s multiple comparisons test). Stress alone does not enhance freezing level because immediate shock alone cannot enhance freezing level in the behavioral experiment in the previous result. \**p*< 0.05, \*\**p*< 0.01 Data are mean ± s.e.m.

**EV 3.**
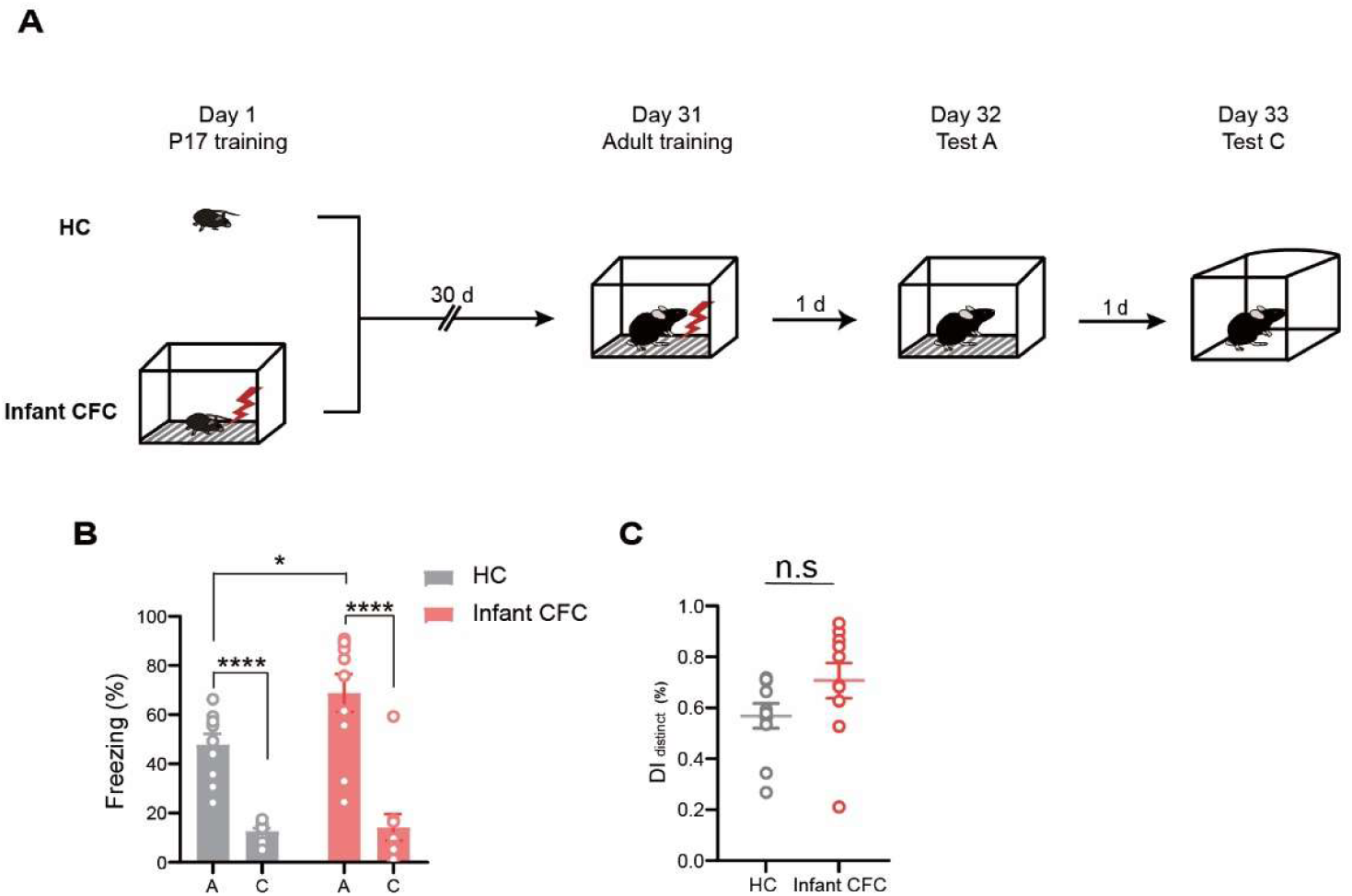
Enhancement of adult fear memory does not lead to memory generalization. **(A)** Behavioral paradigm for context-generalization test (n=10 mice per group). Mice were stayed in the home cage or underwent CFC. Thirty days later, all mice underwent a second CFC session. Freezing behavior was assessed in the conditioned context 24 h later and in a distinct context 24 h thereafter. **(B)** There was significant difference between Context A and Context C in freezing levels (*p*<0.0001, paired t-test). **(C)** There was no significant different between two groups in discrimination index (DI, *p*=0.1166, unpaired t-test). n.s, not significant. \**p*<0.05, \*\*\*\**p*< 0.0001. Data are mean ± s.e.m

**EV 4.**
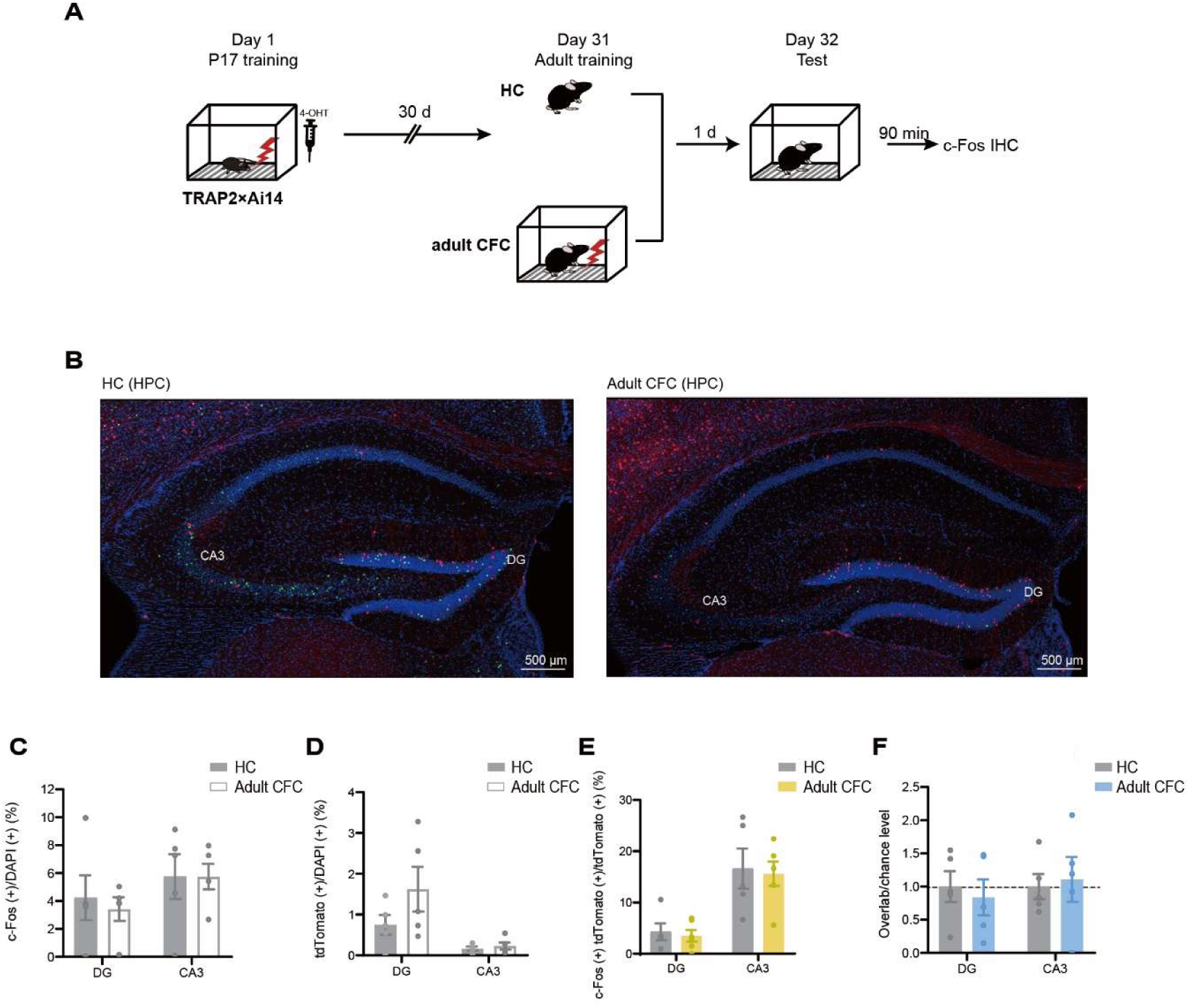
The engram formed in infancy is not recruited hippocampus (HPC) in adulthood. **(A)** Behavioral paradigm **(B)** Representative confocal images. Scale bar: 500 μm and 20 μm, **(C-F),** Quantification of cFos (+) cells, tdTomato (+) cells, their overlap, and chance-normalized overlap in the hippocampus (HPC). There were no significantly difference in cFos (+) cells, in tdTomato (+) cells, their overlap, and chance-normalized overlap. n.s, not significant. Data are mean ± s.e.m

